# Transposable elements in the genome of the lichen-forming fungus *Umbilicaria pustulata*, and their distribution in different climate zones along elevation

**DOI:** 10.1101/2021.06.24.448634

**Authors:** Francesco Dal Grande, Veronique Jamilloux, Nathalie Choisne, Anjuli Calchera, Malte Petersen, Meike Schulz, Maria A. Nilsson, Imke Schmitt

## Abstract

**Background:** Transposable elements (TEs) are an important source of genome plasticity across the tree of life. Accumulating evidence suggests that TEs may not be randomly distributed in the genome. Drift and natural selection are important forces shaping TE distribution and accumulation, acting directly on the TE element or indirectly on the host species. Fungi, with their multifaceted phenotypic diversity and relatively small genome size, are ideal models to study the role of TEs in genome evolution and their impact on the host’s ecological and life history traits. Here we present an account of all TEs found in a high-quality reference genome of the lichen-forming fungus *Umbilicaria pustulata*, a macrolichen species comprising two climatic ecotypes: Mediterranean and cold-temperate. We trace the occurrence of the newly identified TEs in populations along three replicated elevation gradients using a Pool-Seq approach, to identify TE insertions of potential adaptive significance.

**Results:** We found that TEs cover 21.26 % of the 32.9 Mbp genome, with LTR Gypsy and Copia clades being the most common TEs. Out of a total of 182 TE copies we identified 28 insertions displaying consistent insertion frequency differences between the two host ecotypes across the elevation gradients. Most of the highly differentiated insertions were located near genes, indicating a putative function.

**Conclusions:** This pioneering study into the content and climate niche-specific distribution of TEs in a lichen-forming fungus contributes to understanding the roles of TEs in fungal evolution. Particularly, it may serve as a foundation for assessing the impact of TE dynamics on fungal adaptation to the abiotic environment, and the impact of TE activity on the evolution and maintenance of a symbiotic lifestyle.

## Background

Transposable elements (TEs) are DNA sequences that self-propagate across genomes (1). TEs are a ubiquitous component of almost all prokaryotic (2) and eukaryotic genomes such as plants (e.g., (3,4), fungi (5) and animals (6,7)). Eukaryotic TEs fall into two broad classes: DNA transposons that use a cut-and-paste mechanism for their transposition, and retrotransposons, that move via a reverse transcribed RNA intermediate via a copy-and-paste mechanism. TEs can be further classified into superfamilies and families based on specific sequence features (8–10). Most TEs present in eukaryotic genomes are genomic fossils, i.e. inactive remnants of once active copies (11,12). Their variation in copy number and size is responsible for much of the large differences in genome size observed even among closely related species (13–15). On the other hand, the most recent, likely active, transposable fraction of the repeatome – all repeated sequences except microsatellites – remains silenced under normal conditions. TEs are activated by ontogenetic factors and/or environmental cues (16,17). By their repetitive nature TEs provide hotspots for ectopic (non-homologous) recombination and induce chromosomal rearrangements as well as gene shuffling leading to loss of genomic portions or expansion of gene copy numbers. Being mobile, TEs can further locate in coding or regulatory regions, thus strongly affecting gene expression and gene structure and/or function. TEs can thus passively and actively impact genome plasticity, and extensively shape eukaryotic genome evolution (18,19).

TEs generate evolutionary novelty and respond to environmental change, indicating that they are likely to play a relevant role in adaptation (20–26). The relationship between TEs and environmental adaptation is complex, as both activation and repression of transposition in response to environmental changes have been reported (27–29). Most TEs remain silent and evolve in a neutral fashion, while only a minor fraction has adaptive roles (e.g., (30)). Several studies have suggested that the presence of a certain number of potentially active TEs may increase the genome’s ability to cope with environmental stress in a variety of ways, e.g. via major genomic rearrangements (31), TE-driven creation of new regulatory networks involving genes in the TEs’ proximity (32–35), and/or genome alteration via newly generated TE copies (36). As such, TEs can be a major source of intra-population genetic variation in response to environmental pressures (e.g., (37,38)). For instance, TE composition and/or copy number variation in response to micro-climatic conditions was reported for natural populations of wild barley, *Arabidopsis thaliana* (10,39), *A. arenosa* (40), and several *Brassicaceae* species (41). However, there is a general lack of understanding on how environment influences TE abundance and the activity of most TEs in most non-model species. The range and phenotypic consequences of the heritable mutations produced through TE mobilization remain largely unknown.

Fungi are a diverse group of organisms colonizing all habitats on Earth. Their remarkable diversity in terms of morphologies, life-styles, genome sizes, reproductive modes, and ecological niches makes them an ideal group for comparative genomics. Due to their relatively small genome size compared to plants and animals (e.g., 37 Mbp on average in Ascomycota and 46 Mbp in Basidiomycota; (42)), fungal genomes are easier to assemble and annotate. The past decade has seen an extraordinary increase in fungal genomic research, also in the area of TE research. The increased availability of high quality assemblies for a large numbers of fungi has enabled kingdom-wide comparative studies (5,43). The TE content of fungal genomes is variable, typically ranging from 0 to 30%, with up to 90% in the plant-pathogen *Blumeria graminis* (44,45). Retrotransposons with long terminal repeats (LTR) are the most abundant TE elements in fungal genomes. Several studies have shown that TEs are a major driving force for adaptive genome evolution in fungi (46), especially in fungal plant pathogens (43,47). In fact, animal-related and pathogenic fungi tend to have more TEs inserted into genes than fungi with other lifestyles, and may play an important role in effector gene diversification (48,49). Furthermore, TE content in fungi seems to be correlated with the mode of reproduction, with sexual fungi displaying a higher TE load (50). Surprisingly, lichen-forming fungi, a group of highly diverse, ecologically obligate biotrophs, have been more or less completely neglected in TE research. Lichens are textbook examples of ecologically successful symbioses being the result of a tightly integrated relationship between a fungus, typically an ascomycete, and green algae and/or cyanobacteria (51). Lichens, due to their ability to tolerate environmental extremes, their specialized nutritional mode involving more or less strictly selected photosynthetic symbionts, and their varied morphologies and modes of reproduction represent an important missing piece of the puzzle in our attempt to understand the impact of TE activity on the evolutionary trajectory and architecture of fungal genomes.

Here we provide the first in-depth report on the abundance and distribution of TEs in the genome of a lichen-forming fungus, the ascomycete *Umbilicaria pustulata*. *U. pustulata* is a widespread macrolichen that grows attached to rocks from southern Europe to northern Scandinavia. Population genomics analyses revealed the presence of otherwise morphologically indistinguishable ecotypes in *U. pustulata*, i.e. intra-specific lineages, differentially adapted to the Mediterranean and cold-temperate climate zone, and interacting with different algal symbiont communities (52,53). The availability of a high-quality, PacBio-based reference assembly (54), together with marked genome-wide climatic niche differentiation data (52), and the possibility to sample this widespread and abundant species along replicated elevation gradients make *U. pustulata* an ideal model to study the TE content of a lichen-forming fungal genome and its potential link to intra-specific adaptive variation. Specifically, we asked the questions: i) How diverse is the repeatome in *U. pustulata*?; ii) To what extent does TE abundance vary between populations and across gradients?; iii) Are there ecotype-specific TE insertions, and if so, where are they located? To address these questions, we tracked the insertion frequencies of the newly annotated TEs in populations representing the Mediterranean and the cold-temperate ecotypes of the species. To disentangle general trends from local differentiation, we sampled populations across three elevational gradients each encompassing the Mediterranean and the cold-temperate climate zone.

## Results

### *TE landscape in* U. pustulata

The repeatome spans 21.26 % of the *U. pustulata* genome length (Supplementary Table 1). We annotated 119 TE consensus sequences for a total of 5,956 TE copies (704 of which full-length), 6,758 TE fragments, for a cumulative coverage of 6,996,427 bp (Table 2, Supplementary Tables 1, 2). Retrotransposons (Class I) cover 15.6% of the genome of *U. pustulata*, while DNA transposons (Class II) cover 3.5%. Among the Class I elements, Gypsy are the most represented (8.8% of the genome), followed by Copia elements (4.1%). Helitron are the most abundant elements within the Class II (1.7%), followed by Terminal Inverted Repeats (TIR; 1.2%).

TE copies have a median nucleotide identity of ~90% with their respective TE family consensus sequence, ranging from 88.7% of Helitron (Class II) and 86.2% of LTR elements (Class I) to 95.3% for PiggyBac (Class II) and 94% for LINE elements (Class I). The distribution of TE copy identity to their family consensus sequences suggests recent activity (Fig. 1, Supplementary Table 3).

**Fig. 1.**
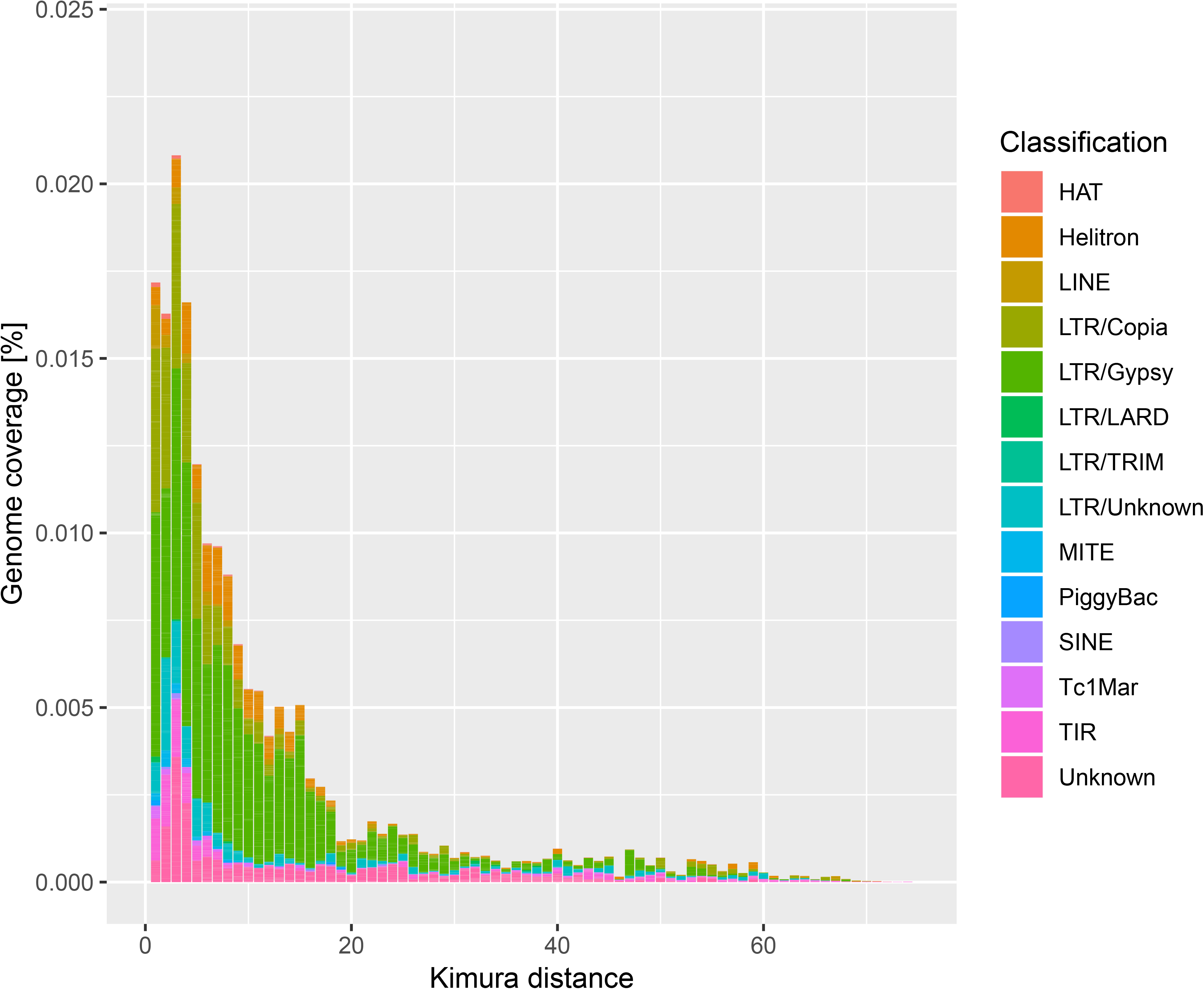
Repeat landscape plot in *U. pustulata*. Sequence divergence of each TE copy from the corresponding consensus sequence was measured based on the Kimura (K2P) distance method. The further to the left a peak in the distribution, the younger the corresponding TE fraction generally is.

### *TE variation across* U. pustulata *populations*

We used the PoPoolationTE2 pipeline (55) on the *U. pustulata* reference genome (54) to detect variations in TE frequencies in 15 natural populations across three replicated elevational gradients.

After manual curation we retained 182 TE loci belonging to 12 superfamilies with a minimum physical coverage of 16 (Table 3A, Supplementary Table 4). Of these, 68 insertions were fixed across populations, i.e., they had a minimum frequency of 0.95 within each population. Copia elements were the most frequently detected loci, representing 43% (49 loci) of all polymorphic insertions, followed by TIR elements (19.3%, 22 loci) (Table 3B).

**Table 1.** Populations ID, coordinates, elevations and Pool-Seq read number for 15 *U. pustulata* populations along three elevational gradients.

| Country | Population ID | Lat | Long | Elevation<br>m a.s.l. | Paired-end read # | mean read<br>length |
| --- | --- | --- | --- | --- | --- | --- |
| Italy | IT1 | 40,7577 | 9,0794 | 176 | 29162770 | 99,3 |
|  | IT2 | 40,7778 | 9,0546 | 297 | 28279628 | 99,3 |
|  | IT3 | 40,8503 | 9,1119 | 588 | 26570943 | 99,4 |
|  | IT4 | 40,8568 | 9,134 | 842 | 31720828 | 99,4 |
|  | IT5 | 40,8573 | 9,1642 | 1125 | 31755901 | 99,4 |
|  | IT6 | 40,8524 | 9,1732 | 1303 | 32064853 | 99,4 |
| Spain 1 | ESii1 | 40,2028 | -5,2334 | 706 | 26758269 | 141,8 |
|  | ESii2 | 40,2069 | -5,2327 | 887 | 24295101 | 141,7 |
|  | ESii3 | 40,2116 | -5,2337 | 1082 | 29236274 | 141,9 |
|  | ESii4 | 40,2183 | -5,2335 | 1258 | 33333561 | 141,6 |
|  | ESii5 | 40,2253 | -5,2375 | 1480 | 24672545 | 141,7 |
|  | ESii6 | 40,2322 | -5,2389 | 1699 | 26690508 | 141,5 |
| Spain 2 | ESi1 | 39,9946 | -4,8679 | 477 | 28862057 | 99,5 |
|  | ESi2 | 40,2899 | -4,9927 | 859 | 37303042 | 99,5 |
|  | ESi3 | 40,323 | -5,0173 | 1417 | 35351050 | 99,5 |

**Table 2A.** Summary of class I and II TE elements found in the *U. pustulata* genome.

| class | Total length | no. copies | no. full length copies | median identity* | median length |
| --- | --- | --- | --- | --- | --- |
| class II | 1146170 | 1863 | 156 | 91,4 | 657,9 |
| class I | 5118614 | 2902 | 465 | 90,3 | 1162,5 |
| Unknown | 731643 | 1191 | 83 | 88,1 | 323,4 |

**Table 2B.** Summary of TE elements subdivided into superfamilies for the *U. pustulata* genome.

| class | order | superfamily | no. elements | total length | no. copies | no. full length copies | median identity* | median length |
| --- | --- | --- | --- | --- | --- | --- | --- | --- |
| class II | DHX | Helitron_01 | 7 | 553513 | 680 | 23 | 88,7 | 498,6 |
|  | DTA | HAT | 1 | 24206 | 80 | 4 | 89,98 | 186,5 |
|  | DTB | PiggyBac | 1 | 12236 | 10 | 4 | 95,3 | 1481,0 |
|  | DTT | Tc1Mar | 4 | 104574 | 139 | 28 | 89,6 | 1029,4 |
|  | DTX | TIR | 18 | 380415 | 824 | 86 | 92,0 | 648,2 |
|  | DXX | MITE | 4 | 71226 | 130 | 11 | 93,0 | 521,0 |
| class I | RII | LINE | 5 | 317234 | 155 | 33 | 94,0 | 923,1 |
|  | RLC | Copia | 25 | 1333809 | 865 | 166 | 92,0 | 1350,2 |
|  | RLG | Gypsy | 23 | 2904582 | 1296 | 215 | 89,8 | 1246,0 |
|  | RLX | LTR | 15 | 538504 | 550 | 46 | 86,2 | 942,6 |
|  | RXX | LARD | 1 | 20415 | 25 | 1 | 816,6 | 383 |
|  | RXX | TRIM | 1 | 4070 | 11 | 4 | 96,8 | 126 |
|  | No | Unknown | 14 | 731643 | 1191 | 83 | 88,1 | 323,4 |
| *total* |  |  | *119* | *6996427* | *5956* | *704* | *147,1* | *743,0* |

\*Identity= % sequence similarity between TE copy and the respective consensus sequence

**Table 3A.** TE copy insertion in 15 populations of *U. pustulata* (min. physical coverage: 16x).

| TE family | copy no. | % |
| --- | --- | --- |
| Copia | 62 | 34,1 |
| TIR | 31 | 17,0 |
| Unknown | 23 | 12,6 |
| Helitron | 22 | 12,1 |
| Gypsy | 16 | 8,8 |
| LTR | 10 | 5,5 |
| MITE | 8 | 4,4 |
| LARD | 5 | 2,7 |
| TC1Mar | 2 | 1,1 |
| HAT | 1 | 0,5 |
| LINE | 1 | 0,5 |
| Piggybac | 1 | 0,5 |

**Table 3B.**
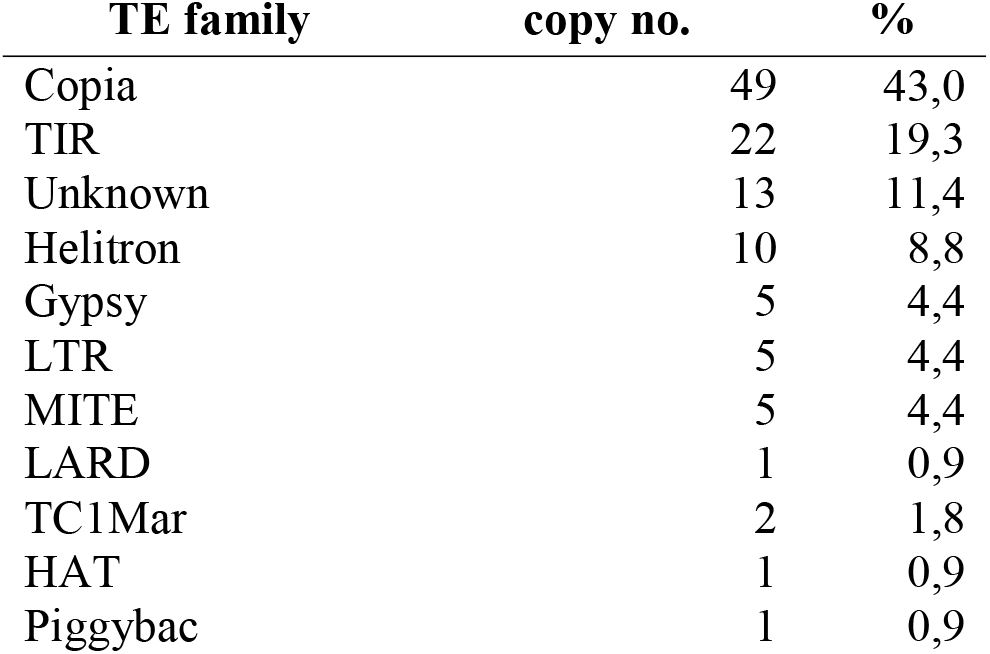
Polymorphic TE copy insertion in populations.

We further compared population structure based on 447,470 genome-wide SNPs (dataset available at: https://doi.org/10.6084/m9.figshare.14784579) with the population divergence based on the variations of TE frequencies across populations. Both SNP-based and TE frequency-based ordinations show that populations can be grouped into two clearly distinct clusters, corresponding to the Mediterranean and cold-temperate ecotypes of the lichen-forming fungus *sensu* Dal Grande et al. (2017) (52) (Fig. 2).

**Fig. 2.**
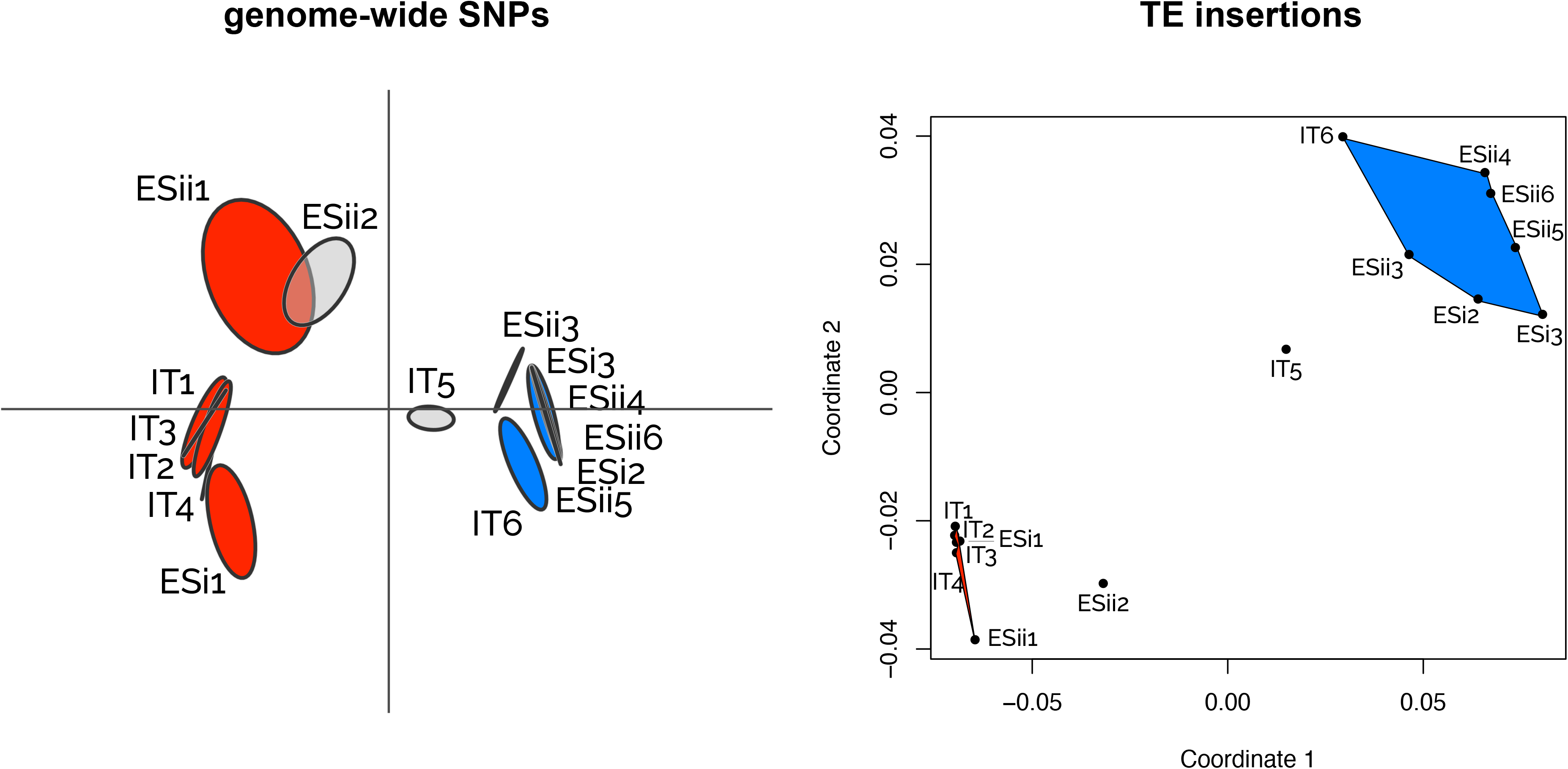
Left: Pattern of genetic structure among populations based on pairwise F_ST_ genetic distances calculated on 447.470 polymorphic SNPs. Right: Non-metric multidimensional scaling (NMDS) ordination plot illustrating population structure based on TE copy insertion frequencies in 15 populations of *U. pustulata*. IT: Italian gradient, ES: Spanish gradients (i, ii). The populations from Mediterranean climate (red) and cold temperate climate (blue) form clusters (with the exception of IT5 and ESii2 which have an intermediate position).

### Variations of TE frequencies between ecotypes

We identified TE loci that were highly differentiated (hdTEs) between the two ecotypes, because these loci might represent differential fixation/loss between ecotypes and have particular functional relevance. We identified 28 hdTEs (Table 3C). Of these, seven were exclusively found in the cold-temperate populations, 19 showed significantly higher frequency in the cold-temperate populations, and one was more abundant in the Mediterranean populations (a short Copia11 fragment in scaffold9_123163). One Copia element was almost exclusively found in the two Spanish gradients (an almost full-length Copia11 copy in scaffold9_1443709). This insertion was absent in the Mediterranean climatic zone and linearly increased in abundance with elevation.

**Table 3C.**
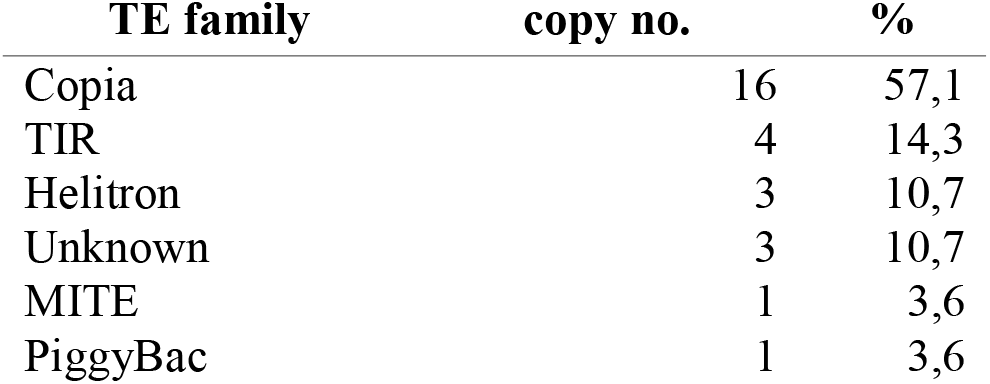
hdTEs between *U. pustulata* ecotypes.

| TE family | copy no. | % |
| --- | --- | --- |
| Copia | 16 | 57,1 |
| TIR | 4 | 14,3 |
| Helitron | 3 | 10,7 |
| Unknown | 3 | 10,7 |
| MITE | 1 | 3,6 |
| PiggyBac | 1 | 3,6 |

The analysis of hdTEs between ecotypes showed an overrepresentation of Copia elements (16 loci, 57.1%). Among hdTEs we also found 4 TIR, 3 Helitron, 3 unknown, 1 MITE and 1 PiggyBac element. Compared to all other TE insertions detected across populations, hdTEs were significantly more similar to their consensus sequence (Wilcoxon signed rank sum test p < 0.0001 both in terms of sequence identity and length coverage). Eighteen hdTEs displayed sequence identity and length coverage towards their respective consensus sequence greater than 95%.

### Potential functional impact of TE insertions

One hundred and three out of 114 polymorphic TE loci were inserted either inside a gene (27 TE loci, 25 in coding positions) or in a possible regulatory region (in the 1-kb region surrounding a gene). These include all except two hdTEs (Supplementary Table 3).

## Discussion

### *The* U. pustulata *repeatome*

In this work we studied the content of transposable elements in the genome of the lichen-forming fungus *U. pustulata*. Furthermore we analyzed the variation in TE insertion frequency in populations representing two ecotypes distributed along three gradients spanning the elevational range of the species, i.e. from the Mediterranean to cold-temperate climate zones.

The repeat content in *U. pustulata* of 21% is rather high, compared to the repetitive content in other fungal genomes, which typically ranges from 0 to 30% (56,57). It is also higher than the predicted 15% TE content in another lichen-forming fungus, the Eurotiomycete *Endocarpon pusillum* (58). The *U. pustulata* TE landscape is particularly rich in retrotransposons (class I), especially the LTR retrotransposons Gypsy and Copia. This is a general feature in fungi. The class I/class II genomic coverage ratio of 1.56 is in line with what has been reported for Ascomycetes (0.78-4.23; (57)).

A substantial portion of the annotated TE copies are highly similar to their consensus, which is often interpreted as a signature of rapid and recent bursts of TE activity in the genome (e.g., (59)). Some TE families, such as Gypsy, on the other hand, displayed a broader range of identity rate with their consensus, suggesting slower colonization of the *U. pustulata* genome with these elements. In the absence of a molecular clock for *U. pustulata* it is, however, difficult to precisely evaluate the time when the TE bursts possibly occurred, and how much time it took for the TEs to spread in the genome.

Population-level analyses of TE insertion frequencies in 15 populations of *U. pustulata* along three elevational gradients showed that a substantial part of the TEs can be considered as stable and fixed among populations. The clustering of populations based on the detected TE loci across the three gradients recapitulated almost exactly the population divergence based on genome-wide SNPs. This suggests that TE variation is mainly a result of drift between populations. The predominant evolutionary neutrality of TE variation has already been reported for other groups of organisms, such as nematodes (60), and other fungi (61).

### Ecotypic differentiation patterns of TE insertions and their potential functional impact

Although adaptive TE insertions may be marginal compared to the overall repeatome dynamics (61), it is broadly recognized that TEs can play important regulatory roles and may contribute substantially to adaptive evolution in a variety of organisms (25,27,62,63). To identify TE insertions likely linked to climatic niche we studied loci where the TE frequencies were significantly differentiated by fungal ecotype, recurrently across the gradients (hdTEs). Overall, the high similarity of hdTEs to their consensuses, the high variability in insertion frequency among populations – often linearly correlated with elevation – as well as the presence of gradient-specific insertions suggest that most of the hdTEs have recently been active in *U. pustulata* and are possibly still active, in particular in populations located in the cold-temperate climate zone.

Copia retrotransposons are the younger, most active elements of the *U. pustulata* repeatome. When Copia elements are in proximity of a gene, their regulatory role is typically exerted via regulation of gene expression by small RNAs, whereas when inserted within genes they can give rise to alternative splice variants (39,64). Genome expansion related to retrotransposon amplification has been shown to occur in plants as a result of environmental adaptation (e.g., (65,66)). Global transcriptomic responses of Copia elements have been linked to heat stress in *Arabidopsis* spp. (41) and to various environmental stresses in *Eucalyptus* (67).

The identified hdTEs are prime candidates for future functional validation, e.g. via targeted transcriptomic and proteomic analyses, to test whether and how they influence adaptation of the lichen ecotype to different climatic niches. Particularly interesting in this regard could be the effects of TEs inserted near i) *genes involved in cell wall biosynthesis*: a Copia element near a putative GPI ethanolamine phosphate gene, controlling membrane-to-cell wall transfer of fungal adhesins by membrane-anchored transglycosidases (68); a TIR element near *Sac7*, a known activator of the small GTPase *RHO1*, which plays an essential role in the control of cell wall synthesis and organization of the actin cytoskeleton (69); ii) *genes involved in nutrient assimilation*: a Copia element near a NADP-specific glutamate dehydrogenase, a key enzyme in the assimilation of alternative nitrogen sources through ammonium (70); an Helitron element near an acid protease, whose secretion grants access to the carbon and mineral nutrients within proteins in the cells of the plant host in fungal endophytes (71); an unknown TE element inserted near an inositol-pentakisphosphate 2-kinase, an enzyme involved in the decomposition of organic phosphates, whose activity is modulated by environmental pH (72); iii) *genes involved in DNA repair mechanisms*: a Copia element near a putative DNA glycosylase, a gene involved in single-base excision repair mechanisms (73); iv) *genes involved in reproduction and environmental sensing*: an unknown TE element located near a conidiation-specific gene, which plays a role in balancing asexual and sexual development, a process regulated by several factors including light, temperature, humidity, and nutrient availability (74,75); v) *genes involved in secondary metabolism*: a PiggyBac element within a type-I polyketide gene cluster containing fixed nonsense mutations in its core biosynthetic gene only in the cold-temperate climate zone (76). TEs have been previously identified as regulators of biosynthetic gene clusters in ascomycetes: the lower expression of the penicillin cluster in *Aspergillus nidulans* in the absence of *Pbla* element is a typical example (77).

### Outlook and future perspectives

To our knowledge, this is the first in-depth report on a lichen repeatome, based on a highly contiguous and complete PacBio-based reference assembly. As more consensus TE libraries will become available in the future, as a result of improved sequencing and assembling technologies, the study of the repeatome of lichen-forming fungi will contribute key insights to the understanding of TE evolution, in particular in the following research areas:

1. *Role of reproductive mode on TE abundance and composition*: the dynamics in TE load according to the reproductive modes are still a matter of debate. Theoretically sexual reproduction may either facilitate TE accumulation by providing a means of spreading to all individuals in a population, or restrain TE accumulation via purifying selection (50). On the other hand, TE movements may constitute an important source of genome plasticity compatible with adaptive evolution in predominantly asexual species (60). Broad-scale comparative analyses of different sexual and asexual lineages in both nematodes and arthropods revealed no evidence for differences in TE load according to the reproductive modes (78,79). In fungi, however, a recent study suggests that sex might be responsible for the evolutionary success of TEs, by showing that TE loads decrease rapidly under asexual reproduction (50). Lichens are ideal study systems to address this question as congeneric, closely-related species often differ strikingly in their modes of reproduction (80,81). In our case, the sister species of the predominantly asexual *U. pustulata, U. hispanica,* reproduces mainly via sexual ascospores (82,83).
2. *Link between TE content and fungal life strategies*: TE count tends to be elevated in fungal plant symbionts (84). This is because recurrent adaptation to symbiosis seems to involve relaxed genome control against duplications, TE proliferation and overall growth in genome size (63). About half of the currently described ascomycete species are involved in a lichen symbotic association. This symbiotic lifestyle is believed to have arisen independently on several occasions in the evolution of Ascomycota (51). Comparing the repeatome of several unrelated lichen-forming fungi across the Fungi will provide important basal information to understand the evolutionary consequences of the symbiotic lifestyle on the fungal repeatome.
3. *Intra-specific variation and role of TEs in adaptive evolution*: several studies have shown that TE insertion patterns may differ between closely related fungal species occupying different niches (e.g., *Ustilago maydis* and *Sporisorium scitamineum,* (85)) or even between strains within the same species (*Magnaporthe grisea*, (86)). Many lichen species are characterized by wide ecological amplitudes, with distributional ranges spanning multiple climate zones. Furthermore, long-lived, sessile organisms such as lichens are more likely to experience strong selective pressures resulting in particularly abrupt genetic breaks between differentially selected populations over short distances (52,87). Lichens are therefore ideal systems to test the intra-specific differentiation in TE content and its potential role in affecting host fitness in different environments.
4. *TE content in lichen-associated photobionts*: Nearly 40 genera of green algae (~100 species) have been reported from lichen symbioses. Studies on the TE content of green algae are scarce. While the TE abundance seems to be low in the green algal lineage (88,89), TEs may have important functional roles. For instance, TEs may have considerably contributed for gene regulatory sequences evolution in the green algal model species *Chlamydomonas reinhardtii* (89). TEs were reported as the major driver of chromosome specialization in two out of the 20 chromosomes in the marine algal *Ostreococcus tauri*, the smallest free-living eukaryote, possibly contributing to environmental niche adaptation and modulation of reproduction (90). Lichen photobionts are an interesting and highly diverse group of unicellular eukaryotes to study in relation to TE diversity and evolution, especially considering the high symbiotic specificity, the high intra-specific diversity and strong environmental structuring found in many taxa (91–95).

In summary, our pioneering study into TE content and variation of a lichen-forming fungus provides valuable baseline data for future investigations. It opens up new perspectives for targeted analyses of the potential effect of TE dynamics on the evolution, fitness and adaptability of *U. pustulata*, and more generally of lichen-forming fungi, and other symbiotic systems.

## Methods

### *The genome of* U. pustulata

We used the genome assembly by Greshake Tsovaras et al. (54) as reference for TE prediction and annotation (accession VXIT01000000, BioProject: PRJNA464168). The haploid genome of *U. pustulata* is 32.9 Mbp long, with 43 scaffolds, and an N50 length of >1.8 Mbp.

### *Pool-Seq sequencing of 15* U. pustulata *populations*

To predict the copy insertion frequencies at TE loci across three elevational gradients, we used whole-genome sequencing data from pools of individuals from 15 natural lichen populations (100 lichen thalli per population). The 15 pools were collected along three elevational gradients in Southern Europe, i.e. Mount Limbara (Sardinia, Italy; 6 populations, IT), Sierra de Gredos (Sistema Central, Spain; 6 populations, ESii) and Talavera-Puerto de Pico (Sistema Central, Spain; 3 populations, ESi) (Table 1), as described in (96). Individuals were pooled in equimolar concentrations and each pool was sequenced on an Illumina HiSeq platform (2 x 100 bp for IT and ESi, 2 x 150 bp for ESii). The Pool-Seq data was quality-filtered using Trimmomatic v0.39 (97) with a length cutoff of 80 bp and a quality cutoff of 26 in a window of 5 bp. Reads with N’s were removed and an additional quality trimming using a modified Mott algorithm was performed using the script *trim-fastq.pl* from the PoPoolation v1.2.2 pipeline (98). After trimming, the sequencing depth varied between 24.3 and 37.3 million paired-end reads (Table 1).

### *De novo TE prediction: building a* U. pustulata *TE-consensus library*

We used the TEdenovo pipeline from the REPET package v2.5 (99,100) to generate a TE-consensus library in *U. pustulata*. Briefly, the pipeline was used to perform a self-alignment of the reference genome to detect repeats, to cluster the repetitions, and to perform multiple alignments from the clustered repetitions to create consensus TE sequences. Consensus TEs were subsequently classified using the PASTEClassifier pipeline v2.0 (101), which follows Wicker’s classification (8) using structural and homology-based information (i.e., terminal repeats, poly(A) tails, ORFs, tandem repeats, etc.) and the following databases: ‘repbase20.05_ntSeq_cleaned_TE.fa’, ‘repbase20.05_aaSeq_cleaned_TE.fa’ and ‘ProfilesBankForREPET_Pfam27.0_GypsyDB.hmm’ (https://urgi.versailles.inra.fr/download/repet). We set the *minNbSeqPerGroup* parameter to 3 (i.e., 2n+1) because *U. pustulata* is haploid. All remaining parameters used for these analyses can be found in the TEdenovo and TEannot configuration files (Additional Files 1, 2).

We then performed extensive automated as well as manual curation of the TE consensus library to minimize redundancy as well as false positives. For this purpose, we first performed a two-step annotation (102) on contigs longer than 5 kbp, i.e. 1^st^ round: steps 1 - taking all matches found by BLASTER, RepeatMasker and CENSOR, 2 - normal and random, 3 - using Grouper, Recon and Piler as clustering methods, 7 - removing duplicated/spurious fragments and applying the long join procedure for nested copies of TEs identified by the TEannot pipeline part. We only retained TE consensus sequences having at least one Full-Length Copy (FLC; i.e. length of fragments between 95% and 105% of consensus length) to build the final TE library. This was followed by a 2^nd^ round consisting of TEannot steps 1, 2, 3, 4, 5, 7 and 8 using the final TE library to annotate the genome.

Finally we performed a copy-divergence analysis of TE classes, based on Kimura distances by calculating Kimura 2-parameter divergence (103) between each TE copy and its consensus sequence using the utility scripts provided in the RepeatMasker package. These were also used to construct a TE landscape divergence plot by grouping copies within TE superfamilies and calculating the percentage of the genome occupied by each TE superfamily.

### *Evaluation of TE copy insertion frequencies across the different* U. pustulata *populations*

We used the PoPoolationTE2 v1.10.04 pipeline (55) to compute population-wide TE copy insertion frequencies of the curated TE library across the 15 populations described above. For this, we performed a ‘joint’ analysis using both quantitative and qualitative information extracted from paired-end reads mapping on the TE-annotated reference genome and a set of reference TEs to detect TE copy insertion frequencies in populations. Frequency values in this case correspond to the proportion of individuals in a population for which a TE copy is present at a given locus.

We used the curated *U. pustulata* TE library and the *U. pustulata* reference genome described above to produce the ‘TE-merged’ reference file (available at: https://doi.org/10.6084/m9.figshare.14784579) and the ‘TE-hierarchy’ file (Additional File 3) as follows. Sequences corresponding to the TE annotations were extracted and masked in the reference genome using the tools getfasta and maskfasta from the Bedtools suite (104), respectively. The resulting TE sequences were concatenated with the masked genome to form the ‘TE-merged’ reference. For every TE copy we also retrieved TE sequence name, family, and order to build the required ‘TE-hierarchy’ file. For each *U. pustulata* pool, we mapped forward and reverse reads separately against the ‘TE-merged’ reference using the local alignment algorithm BWA-SW v0.7 (105) with default parameters. The obtained SAM alignment files were then converted to BAM files using samtools view v1.9 (106). Paired-end information was restored from the previous alignments using the *se2pe* (--sort) tool from PoPoolationTE2 v1.10.04. Using the *ppileup* tool from PoPoolationTE2 we then created a ppileup file (--map-qual 15) that summarizes, for every base of the genome, the number of PE reads spanning the site – i.e., physical coverage – as well as the structural status inferred from the paired-end reads covering the site (i.e., indicating whether one or both boundaries of a TE insertion are supported by significant physical coverage).

Heterogeneity in physical coverage among populations may lead to discrepancies in TE frequency estimation and in a substantial fraction of sample specific insertion false positives (55). Hence, to reduce the number of false positives, we normalized the physical coverage across the *U. pustulata* populations via a subsampling and a rescaling approach: In order to balance the loss of information with the homogeneity of the TE frequency we used the *stat-coverage* tool from PoPoolationTE2 to obtain information on the physical coverage in our dataset. We then used the *subsamplePpileup* tool (--target-coverage 16) to discard positions with a physical coverage below 16x and rescale the coverage of the remaining sites to that value.

We identified signatures of TE polymorphisms from the previously subsampled file using the *identifySignature* tool following the joint algorithm (--mode joint; --min-count 3; --signature- window minimumSampleMedian; --min-valley minimumSampleMedian). Then, for each identified site, we estimated TE frequencies in each pool using the *frequency* tool. Eventually, we paired up the signatures of TE polymorphisms using *pairupSignatures* tool (--min-distance 100; --max-distance 500), yielding a final list of TE loci in the reference genome with their frequencies for each pool. Each TE insertion was manually checked using IGV v2.5 (107). TE loci predictions with unusually high read coverage, i.e. resulting from spurious alignments to unmasked repeats, were discarded from further analysis. The stringent filters applied here, together with the inability of PoPoolationTE2 to detect nested TEs (55), may lead to an underestimation of TE activity across *U. pustulata* populations. On the positive side, however, such a conservative approach almost certainly eliminates false insertions. TE loci supported by significant physical coverage were considered polymorphic if they had a frequency difference of at least 0.05% among populations. TE loci with frequencies ≥0.95% were considered as fixed in the populations. The similarity of populations based on their TE composition was investigated using nonmetric multidimensional scaling (NMDS) on all detected TE insertion frequencies using the function metaMDS from the vegan package (108) for R (109).

### *Identification of TE loci significantly differentiated between* U. pustulata *ecotypes*

To identify highly differentiated TE loci (hdTEs) between *U. pustulata* ecotypes we performed a differential abundance analysis using the microbiomeSeq (110) and DeSeq2 (111) R packages. For this purpose, we contrasted the normalized relative abundances of all TE copy insertions in DeSeq2 to detect differentially abundant TE copy insertions (at α = 0.01) between populations representing the Mediterranean (populations IT1-4, ESii1, ESi1) and the cold-temperate (IT6, ESii3-6, ESi2-3) ecotypes. From the analysis we excluded populations IT5 and ESii2, because they represent admixed populations of both ecotypes (96).

### Functional characterization

To identify genes potentially impacted by TE insertions, i.e. genes overlapping with TEs or in the proximity of TEs (1 kbp up- or downstream each TE insertion), we cross-referenced the TE annotation file with the gene annotation file (54) using the *intersect* tool of the Bedtools suite (104).

### Population structure based on genome-wide SNPs

Population structure based on genome-wide single-nucleotide polymorphisms (SNPs), i.e. the positional relations among populations based on their genetic distances, was detected by analyzing pairwise quantile distance matrices (0.975, 0.75, 0.5, 0.25, 0.025) based on the pairwise fixation index (F_ST_) among all populations using a three-way generalization of classical multidimensional scaling (DISTATIS; (112)). Briefly, we used the sorted, duplicate-removed BAM files of reads mapped to the *U. pustulata* reference genome. High-quality (i.e. after removing duplicated reads and genomic indels) SNPs were called using SAMtools mpileup and normalized to a uniform coverage of 30 across all populations with PoPoolation2 (113). For this we used the synchronized mpileup file (i.e. ‘sync’ file containing the allele frequencies for every population at every base in the reference genome) and the script *subsample-synchronized.pl* (--without-replacement), excluding positions with a coverage exceeding the 2% of the empirical coverage distribution of each pool. Genetic differentiation (F_ST_) was calculated with *fst-sliding.pl* in PoPoolation2 on the subsampled sync file. We only considered SNPs with a minimum read count of 4 and a minimum mapping quality of 20. A more detailed description of the methods can be found in (96).

## Declarations

### Ethics approval and consent to participate

Not applicable.

### Consent for publication

Not applicable.

### Availability of data and materials

Raw sequences are available in the SRA archive under Bioproject [xxx]. The datasets supporting the conclusions of this article are available in the Figshare repository, https://doi.org/10.6084/m9.figshare.14784579.

### Authors’ contributions

FDG and IS conceived the idea to the study. FDG, VC, NC, AC, MP, and MS analyzed the data. FDG, MP, MN, and IS interpreted the data. FDG produced the figures and wrote the manuscript. All authors read, approved, and commented on the manuscript.

## Acknowledgements

We thank Jürgen Otte (Frankfurt) for laboratory assistance, and Ann-Marie Waldvogel (Cologne) for stimulating discussions during the early phase of this project. Claus Weiland (Frankfurt) provided invaluable support with software installation.

## References

1. Craig N, Chandler M, Gellert M, Lambowitz A, Rice P, SB S. Mobile DNA III. 3rd ed. American Society For Microbiology (ASM), editor. Washington, DC; 2015.

2. Sawyer S, Hartl D. Distribution of transposable elements in prokaryotes. Theor Popul Biol. 1986;30:1–16.

3. Bennetzen JL. Transposable elements, gene creation and genome rearrangement in flowering plants. Curr Opin Genet Dev. 2005;15:621–7.

4. Staton SE, Burke JM. Evolutionary transitions in the Asteraceae coincide with marked shifts in transposable element abundance. BMC Genomics. 2015;16:623.

5. Daboussi M-J, Capy P. Transposable elements in filamentous fungi. Annu Rev Microbiol. 2003;57:275–99.

6. Chalopin D, Naville M, Plard F, Galiana D, Volff J-N. Comparative analysis of transposable elements highlights mobilome diversity and evolution in vertebrates. Genome Biol Evol. 2015;7:567–80.

7. Petersen M, Armisén D, Gibbs RA, Hering L, Khila A, Mayer G, et al. Diversity and evolution of the transposable element repertoire in arthropods with particular reference to insects. BMC Ecol Evol. 2019;19:11.

8. Wicker T, Sabot F, Hua-Van A, Bennetzen JL, Capy P, Chalhoub B, et al. A unified classification system for eukaryotic transposable elements. Nat Rev Genet. 2007;8:973–82.

9. Kapitonov V V., Jurka J. A universal classification of eukaryotic transposable elements implemented in Repbase. Nat Rev Genet. 2008;9:411–2.

10. Quadrana L, Etcheverry M, Gilly A, Caillieux E, Madoui MA, Guy J, et al. Transposition favors the generation of large effect mutations that may facilitate rapid adaption. Nat Commun. 2019;10:1–10.

11. Smit AFA. Interspersed repeats and other mementos of transposable elements in mammalian genomes. Curr Opin Genet Dev. 1999;9:657–63.

12. Lanciano S, Cristofari G. Measuring and interpreting transposable element expression. Nat Rev Genet. 2020;21:721–36.

13. Boulesteix M, Weiss M, Biémont C. Differences in genome size between closely related species: The *Drosophila melanogaster* species subgroup. Mol Biol Evol. 2006;23:162–7.

14. Hawkins JS, Kim HR, Nason JD, Wing RA, Wendel JF. Differential lineage-specific amplification of transposable elements is responsible for genome size variation in *Gossypium*. Genome Res. 2006;16:1252–61.

15. Klein SJ, O’Neill RJ. Transposable elements: genome innovation, chromosome diversity, and centromere conflict. Chromosom Res. 2018;26:5–23.

16. Makarevitch I, Waters AJ, West PT, Stitzer M, Hirsch CN, Ross-Ibarra J, et al. Transposable elements contribute to activation of maize genes in response to abiotic stress. PLOS Genet. 2015;11:e1004915.

17. Rey O, Danchin E, Mirouze M, Loot C, Blanchet S. Adaptation to global change: a transposable element–epigenetics perspective. Trends Ecol Evol. 2016;31:514–26.

18. Feschotte C, Pritham EJ. DNA transposons and the evolution of eukaryotic genomes. Annu Rev Genet. 2007;41:331–68.

19. Pritham EJ. Transposable elements and factors influencing their success in eukaryotes. J Hered. 2009/08/07. 2009;100:648–55.

20. Biémont C, Vieira C. Genetics: Junk DNA as an evolutionary force. Nature. 2006;443:521–4.

21. Schmidt AL, Anderson LM. Repetitive DNA elements as mediators of genomic change in response to environmental cues. Biol Rev. 2006;81:531–43.

22. Oliver KR, Greene WK. Transposable elements: powerful facilitators of evolution. BioEssays. 2009 Jul 1;31:703–14.

23. Hua-Van A, Le Rouzic A, Boutin TS, Filée J, Capy P. The struggle for life of the genome’s selfish architects. Biol Direct. 2011;6:19.

24. Casacuberta E, González J. The impact of transposable elements in environmental adaptation. Mol Ecol. 2013;22:1503–17.

25. Hof AE van’t, Campagne P, Rigden DJ, Yung CJ, Lingley J, Quail MA, et al. The industrial melanism mutation in British peppered moths is a transposable element. Nature. 2016;534:102–5.

26. Schrader L, Schmitz J. The impact of transposable elements in adaptive evolution. Mol Ecol. 2019;28:1537–49.

27. González J, Petrov DA. The adaptive role of transposable elements in the *Drosophila* genome. Gene. 2009;448:124–33.

28. Horváth V, Merenciano M, González J. Revisiting the relationship between transposable elements and the eukaryotic stress response. Trends Genet. 2017;33:832–41.

29. Dubin MJ, Mittelsten Scheid O, Becker C. Transposons: a blessing curse. Curr Opin Plant Biol. 2018;42:23–9.

30. Arkhipova IR. Neutral theory, transposable elements, and eukaryotic genome evolution. Mol Biol Evol. 2018;35:1332–7.

31. Maumus F, Allen AE, Mhiri C, Hu H, Jabbari K, Vardi A, et al. Potential impact of stress activated retrotransposons on genome evolution in a marine diatom. BMC Genomics. 2009;10:1–19.

32. Naito K, Cho E, Yang G, Campbell MA, Yano K, Okumoto Y, et al. Dramatic amplification of a rice transposable element during recent domestication. Proc Natl Acad Sci. 2006;103:17620–5.

33. Guo Y, Levin HL. High-throughput sequencing of retrotransposon integration provides a saturated profile of target activity in *Schizosaccharomyces pombe*. Genome Res. 2010;20:239–48.

34. Ito H, Gaubert H, Bucher E, Mirouze M, Vaillant I, Paszkowski J. An siRNA pathway prevents transgenerational retrotransposition in plants subjected to stress. Nature. 2011;472:115–9.

35. Servant G, Pinson B, Tchalikian-Cosson A, Coulpier F, Lemoine S, Pennetier C, et al. *Tye7* regulates yeast *Ty1* retrotransposon sense and antisense transcription in response to adenylic nucleotides stress. Nucleic Acids Res. 2012;40:5271–82.

36. Dai J, Xie W, Brady TL, Gao J, Voytas DF. Phosphorylation regulates integration of the yeast *Ty5* retrotransposon into heterochromatin. Mol Cell. 2007;27:289–99.

37. Lockton S, Ross-Ibarra J, Gaut BS. Demography and weak selection drive patterns of transposable element diversity in natural populations of *Arabidopsis lyrata*. Proc Natl Acad Sci U S A. 2008;105:13965–70.

38. Stewart C, Kural D, Strömberg MP, Walker JA, Konkel MK, Stütz AM, et al. A comprehensive map of mobile element insertion polymorphisms in humans. PLOS Genet. 2011;7:e1002236.

39. Li ZW, Hou XH, Chen JF, Xu YC, Wu Q, Gonzalez J, et al. Transposable elements contribute to the adaptation of *Arabidopsis thaliana*. Genome Biol Evol. 2018;10:2140–50.

40. Wos G, Choudhury RR, Kolář F, Parisod C. Transcriptional activity of transposable elements along an elevational gradient in *Arabidopsis arenosa*. Mob DNA. 2021;12:1–12.

41. Pietzenuk B, Markus C, Gaubert H, Bagwan N, Merotto A, Bucher E, et al. Recurrent evolution of heat-responsiveness in Brassicaceae COPIA elements. Genome Biol. 2016;17:1–15.

42. Mohanta TK, Bae H. The diversity of fungal genome. Biol Proced Online. 2015;17:8.

43. Lorrain C, Feurtey A, Möller M, Haueisen J, Stukenbrock E. Dynamics of transposable elements in recently diverged fungal pathogens: lineage-specific transposable element content and efficiency of genome defenses. G3 Genes|Genomes|Genetics. 2021;11:jkab068.

44. Cuomo CA, Güldener U, Xu J-R, Trail F, Turgeon BG, Di Pietro A, et al. The *Fusarium graminearum* genome reveals a link between localized polymorphism and pathogen specialization. Science 2007;317:1400–2.

45. Frantzeskakis L, Kracher B, Kusch S, Yoshikawa-Maekawa M, Bauer S, Pedersen C, et al. Signatures of host specialization and a recent transposable element burst in the dynamic one-speed genome of the fungal barley powdery mildew pathogen. BMC Genomics. 2018;19:1–23.

46. Oggenfuss U, Badet T, Wicker T, Hartmann FE, Singh NK, Abraham LN, et al. A population-level invasion by transposable elements triggers genome expansion in a fungal pathogen. bioRxiv. 2021;2020.02.11.944652.

47. Grandaubert J, Lowe RGT, Soyer JL, Schoch CL, Van De Wouw AP, Fudal I, et al. Transposable element-assisted evolution and adaptation to host plant within the *Leptosphaeria maculans*-*Leptosphaeria biglobosa* species complex of fungal pathogens. BMC Genomics. 2014;15:1–27.

48. Fouché S, Plissonneau C, Croll D. The birth and death of effectors in rapidly evolving filamentous pathogen genomes. Curr Opin Microbiol. 2018;46:34–42.

49. Fokkens L, Shahi S, Connolly LR, Stam R, Schmidt SM, Smith KM, et al. The multi-speed genome of *Fusarium oxysporum* reveals association of histone modifications with sequence divergence and footprints of past horizontal chromosome transfer events. bioRxiv. 2018;465070.

50. Bast J, Jaron KS, Schuseil D, Roze D, Schwander T. Asexual reproduction reduces transposable element load in experimental yeast populations. Coop G, Tautz D, Coop G, Charlesworth B, editors. Elife. 2019;8:e48548.

51. Lutzoni F, Pagel M, Reeb V. Major fungal lineages are derived from lichen symbiotic ancestors. Nature. 2001;411:937–40.

52. Dal Grande F, Sharma R, Meiser A, Rolshausen G, Büdel B, Mishra B, et al. Adaptive differentiation coincides with local bioclimatic conditions along an elevational cline in populations of a lichen-forming fungus. BMC Evol Biol. 2017;17:93.

53. Dal Grande F, Rolshausen G, Divakar PKPK, Crespo A, Otte J, Schleuning M, et al. Environment and host identity structure communities of green algal symbionts in lichens. New Phytol. 2018;217:277–89.

54. Greshake Tzovaras BG, Segers FHID, Bicker A, Dal Grande F, Otte J, Anvar SY, et al. What is in *Umbilicaria pustulata*? A metagenomic approach to reconstruct the holo-genome of a lichen. Genome Biol Evol. 2020;12:309–24.

55. Kofler R, Gómez-Sánchez D, Schlötterer C. PoPoolationTE2: Comparative population genomics of transposable elements using Pool-Seq. Mol Biol Evol. 2016;33:2759–64.

56. Cuomo CA, Birren BWBT-M in E. Chapter 34 - The Fungal Genome Initiative and lessons learned from genome sequencing. In: Guide to yeast genetics: functional genomics, proteomics, and other systems analysis. Academic Press; 2010. p. 833–55.

57. Castanera R, López-Varas L, Borgognone A, LaButti K, Lapidus A, Schmutz J, et al. Transposable elements versus the fungal genome: impact on whole-genome architecture and transcriptional profiles. PLoS Genet. 2016;12:1–27.

58. Wang Y-Y, Liu B, Zhang X-LX-Y, Zhou Q, Zhang T, Li H, et al. Genome characteristics reveal the impact of lichenization on lichen-forming fungus *Endocarpon pusillum* Hedwig (Verrucariales, Ascomycota). BMC Genomics. 2014;15:34.

59. Lerat E, Goubert C, Guirao-Rico S, Merenciano M, Dufour AB, Vieira C, et al. Population-specific dynamics and selection patterns of transposable element insertions in European natural populations. Mol Ecol. 2019;28:1506–22.

60. Kozlowski DKL, Hassanaly-Goulamhoussen R, Rocha M Da, Koutsovoulos GD, Bailly-Bechet M, Danchin EGJ. Transposable elements are an evolutionary force shaping genomic plasticity in the parthenogenetic root-knot nematode *Meloidogyne incognita*. Evol Appl. 2021;00:1–23.

61. Muszewska A, Steczkiewicz K, Stepniewska-Dziubinska M, Ginalski K. Transposable elements contribute to fungal genes and impact fungal lifestyle. Sci Rep. 2019;9:1–10.

62. Rebollo R, Romanish MT, Mager DL. Transposable Elements: An abundant and natural source of regulatory sequences for host genes. Annu Rev Genet. 2012;46:21–42.

63. Muszewska A, Steczkiewicz K, Stepniewska-Dziubinska M, Ginalski K. Cut-and-paste transposons in fungi with diverse lifestyles. Genome Biol Evol. 2017;9:3463–77.

64. Bourque G, Burns KH, Gehring M, Gorbunova V, Seluanov A, Hammell M, et al. Ten things you should know about transposable elements. Genome Biol. 2018;19:1–12.

65. Kawakami T, Dhakal P, Katterhenry AN, Heatherington CA, Ungerer MC. Transposable element proliferation and genome expansion are rare in contemporary sunflower hybrid populations despite widespread transcriptional activity of LTR retrotransposons. Genome Biol Evol. 2011;3:156–67.

66. Lee J, Waminal NE, Choi H Il, Perumal S, Lee SC, Nguyen VB, et al. Rapid amplification of four retrotransposon families promoted speciation and genome size expansion in the genus *Panax*. Sci Rep. 2017;7:1–9.

67. Marcon HS, Domingues DS, Silva JC, Borges RJ, Matioli FF, de Mattos Fontes MR, et al. Transcriptionally active LTR retrotransposons in *Eucalyptus* genus are differentially expressed and insertionally polymorphic. BMC Plant Biol. 2015;15:1–16.

68. Essen LO, Vogt MS, Mösch HU. Diversity of GPI-anchored fungal adhesins. Biol Chem. 2020;401:1389–405.

69. Schmidt A, Schmelzle T, Hall MN. The *RHO1*-*GAPs SAC7*, *BEM2* and *BAG7* control distinct *RHO1* functions in *Saccharomyces cerevisiae*. Mol Microbiol. 2002;45:1433–41.

70. Downes DJ, Davis MA, Kreutzberger SD, Taig BL, Todd RB. Regulation of the NADP-glutamate dehydrogenase gene *gdhA* in *Aspergillus nidulans* by the Zn(II)2Cys6 transcription factor LeuB. Microbiol. 2013;159:2467–80.

71. Mayerhofer MS, Fraser E, Kernaghan G. Acid protease production in fungal root endophytes. Mycologia. 2015;107:1–11.

72. Williams SP, Gillaspy GE, Perera IY. Biosynthesis and possible functions of inositol pyrophosphates in plants. Front Plant Sci. 2015;6:67.

73. Krokan HE, Bjørås M. Base excision repair. Cold Spring Harb Perspect Biol. 2013;5:a012583.

74. Fabro G, Alvarez ME. Loss of compatibility might explain resistance of the *Arabidopsis thaliana* accession Te-0 to *Golovinomyces cichoracearum*. BMC Plant Biol. 2012;12:143.

75. Wang Z, Miguel-Rojas C, Lopez-Giraldez F, Yarden O, Trail F, Townsend JP. Metabolism and development during conidial germination in response to a carbon-nitrogen-rich synthetic or a natural source of nutrition in *Neurospora crassa*. MBio. 2019;10:1–17.

76. Singh G, Calchera A, Schulz M, Drechsler M, Bode HB, Schmitt I, Dal Grande F. Climate-specific biosynthetic gene clusters in populations of a lichen-forming fungus. Environ Microbiol. 2021; https://doi.org/10.1111/1462-2920.15605

77. Shaaban M, Palmer JM, El-Naggar WA, El-Sokkary MA, Habib E-SE, Keller NP. Involvement of transposon-like elements in penicillin gene cluster regulation. Fungal Genet Biol. 2010;47:423–32.

78. Bast J, Schaefer I, Schwander T, Maraun M, Scheu S, Kraaijeveld K. No Accumulation of transposable elements in asexual arthropods. Mol Biol Evol. 2016;33:697–706.

79. Szitenberg A, Cha S, Opperman CH, Bird DM, Blaxter ML, Lunt DH. Genetic drift, not life history or RNAi, determine long-term evolution of transposable elements. Genome Biol Evol. 2016;8:2964–78.

80. Poelt J. Das Konzept der Artenpaare bei den Flechten. Vor aus dem Gesamtgebiet der Bot NF Deutsch Bot Ges. 1970;4:187–98.

81. Singh G, Dal Grande F, Cornejo C, Schmitt I, Scheidegger C. Genetic basis of self-incompatibility in the lichen-forming fungus *Lobaria pulmonaria* and skewed frequency distribution of mating-type idiomorphs: implications for conservation. PLoS One. 2012;7:e51402.

82. Sancho LG, Crespo A. *Lasallia hispanica* and related species. Lichenol. 1989;21:45–58.

83. Dal Grande F, Meiser A, Greshake Tzovaras B, Otte J, Ebersberger I, Schmitt I. The draft genome of the lichen-forming fungus *Lasallia hispanica* (Frey) Sancho & A. Crespo. Lichenologist. 2018;50:329–40.

84. Hess J, Skrede I, Wolfe BE, Butti K La, Ohm RA, Grigoriev I V., et al. Transposable element dynamics among asymbiotic and ectomycorrhizal *Amanita* fungi. Genome Biol Evol. 2014;6:1564–78.

85. Dutheil JY, Mannhaupt G, Schweizer G, Sieber CMK, Münsterkötter M, Güldener U, et al. A tale of genome compartmentalization: The evolution of virulence clusters in smut fungi. Genome Biol Evol. 2016;8:681–704.

86. Shirke MD, Mahesh HB, Gowda M. Genome-wide comparison of *Magnaporthe* species reveals a host-specific pattern of secretory proteins and transposable elements. PLoS One. 2016;11:e0162458.

87. Chen JM, Werth S, Sork VL. Comparison of phylogeographical structures of a lichen-forming fungus and its green algal photobiont in western North America. J Biogeogr. 2016;43:932–43.

88. Worden AZ, Lee J-H, Mock T, Rouzé P, Simmons MP, Aerts AL, et al. Green evolution and dynamic adaptations revealed by genomes of the marine picoeukaryotes *Micromonas*. Science. 2009;324:268–72.

89. Philippsen GS, Avaca-Crusca JS, Araujo APU, DeMarco R. Distribution patterns and impact of transposable elements in genes of green algae. Gene. 2016;594:151–9.

90. Derelle E, Ferraz C, Rombauts S, Rouzé P, Worden AZ, Robbens S, et al. Genome analysis of the smallest free-living eukaryote *Ostreococcus tauri* unveils many unique features. Proc Natl Acad Sci U S A. 2006;103:11647–52.

91. Fernández-Mendoza F, Domaschke S, García M a, Jordan P, Martín MP, Printzen C. Population structure of mycobionts and photobionts of the widespread lichen *Cetraria aculeata*. Mol Ecol. 2011;20:1208–32.

92. Widmer I, Dal Grande F, Excoffier L, Holderegger R, Keller C, Mikryukov VSVS, et al. European phylogeography of the epiphytic lichen fungus *Lobaria pulmonaria* and its green algal symbiont. Mol Ecol. 2012;21:5827–44.

93. Dal Grande F, Beck A, Cornejo C, Singh G, Cheenacharoen S, Nelsen MP, et al. Molecular phylogeny and symbiotic selectivity of the green algal genus *Dictyochloropsis* s.l. (Trebouxiophyceae): A polyphyletic and widespread group forming photobiont-mediated guilds in the lichen family Lobariaceae. New Phytol. 2014;202:455–70.

94. Werth S, Sork VL. Ecological specialization in *Trebouxia* (Trebouxiophyceae) photobionts of *Ramalina menziesii* (Ramalinaceae) across six range-covering ecoregions of western North America. Am J Bot. 2014;101:1127–40.

95. Rolshausen G, Dal Grande F, Sadowska-Deś ADAD, Otte J, Schmitt I. Quantifying the climatic niche of symbiont partners in a lichen symbiosis indicates mutualist-mediated niche expansions. Ecography. 2018;41:1380–92.

96. Dal Grande F, Sharma R, Meiser A, Rolshausen G, Büdel B, Mishra B, et al. Adaptive differentiation coincides with local bioclimatic conditions along an elevational cline in populations of a lichen-forming fungus. BMC Evol Biol. 2017;17:93.

97. Bolger AM, Lohse M, Usadel B. Trimmomatic: a flexible trimmer for Illumina sequence data. Bioinformatics. 2014;30:2114–20.

98. Kofler R, Orozco-terWengel P, De Maio N, Pandey RV, Nolte V, Futschik A, et al. PoPoolation: a toolbox for population genetic analysis of next generation sequencing data from pooled individuals. PLoS One. 2011;6:e15925.

99. Quesneville H, Bergman CM, Andrieu O, Autard D, Nouaud D, Ashburner M, et al. Combined evidence annotation of transposable elements in genome sequences. PLoS Comput Biol. 2005;1:0166–75.

100. Flutre T, Duprat E, Feuillet C, Quesneville H. Considering transposable element diversification in de novo annotation approaches. PLoS One. 2011;6:e16526.

101. Hoede C, Arnoux S, Moisset M, Chaumier T, Inizan O, Jamilloux V, et al. PASTEC: An automatic transposable element classification tool. PLoS One. 2014;9:1–6.

102. Jamilloux V, Daron J, Choulet F, Quesneville H. De novo annotation of transposable elements: tackling the fat genome issue. Proc IEEE. 2017;105:474–81.

103. Kimura M. A simple method for estimating evolutionary rates of base substitutions through comparative studies of nucleotide sequences. J Mol Evol. 1980;16:111–20.

104. Quinlan AR, Hall IM. BEDTools: a flexible suite of utilities for comparing genomic features. Bioinformatics. 2010;26:841–2.

105. Li H, Durbin R. Fast and accurate short read alignment with Burrows-Wheeler transform. Bioinformatics. 2009;25:1754–60.

106. Li H, Handsaker B, Wysoker A, Fennell T, Ruan J, Homer N, et al. The sequence alignment/map format and SAMtools. Bioinformatics. 2009;25:2078–9.

107. Thorvaldsdóttir H, Robinson JT, Mesirov JP. Integrative Genomics Viewer (IGV): high-performance genomics data visualization and exploration. Brief Bioinform. 2013;14:178–92.

108. Dixon P. VEGAN, a package of R functions for community ecology. J Veg Sci. 2003;14:927–30.

109. R Core Team. R: A language and environment for statistical computing. Vienna, Austria; 2020.

110. Paulson JN, Stine OC, Bravo HC, Pop M. Differential abundance analysis for microbial marker-gene surveys. Nat Methods. 2013;10:1200–2.

111. Love M, Anders S, Huber W. Differential analysis of count data–the DESeq2 package. Genome Biol. 2014;11:R106.

112. Abdi H, O’Toole AJ, Valentin D, Edelman B. DISTATIS: the analysis of multiple distance matrices. In: 2005 IEEE Computer Society Conference on Computer Vision and Pattern Recognition (CVPR’05) - Workshops. IEEE; 2005.

113. Kofler R, Pandey RV, Schlötterer C. PoPoolation2: identifying differentiation between populations using sequencing of pooled DNA samples (Pool-Seq). Bioinformatics. 2011;27:3435–6.

